# Multi-generational biotic stress increases the rate of spontaneous epimutations in a ROS1-dependent manner

**DOI:** 10.1101/2024.09.19.613880

**Authors:** Zhilin Zhang, Adam Hannan Parker, Aikaterini Symeonidi, Samuel W. Wilkinson, Joost H. M. Stassen, Lisa M. Smith, Jurriaan Ton, Frank Johannes

## Abstract

Mistakes in the maintenance of CG methylation are a source of spontaneous epimutations in plants that can be inherited across generations. The extent to which these stochastic events are affected by prolonged exposure to biotic and abiotic stress remains poorly characterized. Here, we grew Arabidopsis Mutation Accumulation (MA) lines for 12 generations in the presence of two biotic stressors: *Pseudomonas syringae* (Pst) and salicylic acid (SA). We found that multi-generational exposure to Pst and SA led to an 18%-32% and 23%-61% increase in the genome-wide epimutation rate, respectively. These rate increases were mainly targeted to subsets of genes characterized by low steady-state methylation (LM), on average, and a lack of transcriptional responsiveness to stress. We show that these effects are mediated by the DNA demethylase *Repressor of Silencing 1* (ROS1). Loss of ROS1 not only buffers transcriptional responses to biotic stress, but also stabilizes epimutation rates, particularly in LM genes, rendering them insensitive to environmental perturbations. Taken together, our data demonstrates that stress can induce heritable epimutations and highlights a ROS1-mediated link between transcriptional plasticity and DNA methylation maintenance fidelity over generations.

## Introduction

Understanding how plants respond to long-term biotic and abiotic stress is important for plant breeding and for predicting biodiversity loss in the face of climate change. Numerous studies point to histone modifications, small RNAs and DNA methylation as important molecular components in stress responses (1–5). Changes in these epigenetic modifications contribute to the transcriptional adjustments that are necessary to acclimate plant physiology, morphology and growth to suboptimal conditions. These epigenetic changes are typically “reset” to their pre-stress state upon removal of the stressor. In some circumstances, however, an epigenetically-encoded stress memory can be maintained in the absence of the stressor for about one sexual generation (2), as observed for example in temperature, salinity and immune priming (2, 6–9). The maintenance of this memory allows plants and their progeny to react more effectively upon re-exposure to the stressor (2, 8–11). Stress responses, as well as the formation of stress memories, are severely impaired in mutants lacking specific components of the epigenetic machinery (5, 7); hence, epigenetic mechanisms are absolutely required for short-term stress acclimation in plants.

In addition to the above transient epigenetic effects, stress exposure can also elicit more lasting chromatin alterations in the form of “‘spontaneous epimutations” (1, 12, 13). These arise stochastically in plant genomes due to epigenome maintenance failures. Spontaneous epimutations have been studied mainly at the level of DNA methylation (1), where they appear as stochastic methylation gains and losses at individual cytosines or larger clusters of cytosines (12, 14–16). When such events occur at CG dinucleotides they can be stably inherited over thousands of generations (12, 13, 17–19), and may have long-term consequences for plant lineage evolution. Under stress conditions, spontaneous epimutations accumulate much more rapidly in plant genomes than under ambient conditions (6). Studies in Arabidopsis and rice grown for multiple generations under high salinity, drought, or heat stress report a general increase in the number of epimutations over time (6, 20, 21). Whether similar observations hold for a wider range of stressors, particularly biotic stress, remains unknown. An intriguing hypothesis is that the accumulation of epimutations is preferentially targeted to stress-response genes, perhaps as a direct consequence of transcriptional activation (or repression). This “epimutation bias” could generate potentially adaptive epialleles, and rapidly contribute to stress adaptation in the long-term. However, current data on multi-generational stress experiments are rare and this hypothesis has not been tested systematically.

Here we propagated *Arabidopsis thaliana* mutation accumulation (MA) lines for 12 generations with repeated Pst and SA exposure and reanalyzed data of MA lines grown under abiotic stress. We found that multi-generational exposure to biotic or abiotic stress led to a significant increase in the CG epimutation rate, particularly in transcriptionally recalcitrant LM genes. The loss of ROS1 substantially reshaped the epimutation landscape, stabilizing epimutation rates and buffering transcriptional responses during biotic stress exposure. Our data demonstrate that biotic stress induces heritable epimutations and emphasizes a ROS1-mediated connection between transcriptional responses and the maintenance of DNA methylation across generations.

## Results

### Multi-generational biotic stress increases the genome-wide epimutation rate

To evaluate how multi-generational biotic stress affects the accumulation of spontaneous epimutations, we generated *A. thaliana* mutation accumulation (MA) lines in the presence of *Pseudomonas syringae* (Pst) and the biotic stress hormone salicylic acid (SA), along with a control condition (Mock)(18)(**Fig. 1A, Methods**). The MA pedigrees in each treatment consisted of four lineages that were independently propagated for 11 generations (G1 - G11), originating from a common founder plant at G0. Whole-genome bisulfite sequencing (WGBS) measurements were taken at generations G2, G6, and G11 (i.e. in progeny of plants treated for 1, 5 or 10 generations). This resulted in 12 WGBS measurements per treatment (**Table S1**). We found that multi-generational biotic stress significantly increased the genome-wide CG epimutation rate, leading to rapid methylation divergence among MA lineages over generations compared with controls (**Fig. 1B-C, Table S2, Table S3, Methods**). Consistent with previous reports (1, 12, 22), no transgenerational accumulation of epimutations could be detected in sequence context CHG and CHH context (where H = A, T, C) (**Table S4**). Using individual CGs as units of analysis (site-level analysis), we estimated that exposure to Pst led to a 31% increase in the spontaneous methylation gain rate (α) and a 32% increase in the spontaneous methylation loss rate (β), while exposure to SA resulted in a 58% and 63% increase of the gain and loss rates, respectively. These rate increases were highly significant (**Fig. 1B, Table S2, Methods**).

**Fig. 1.**
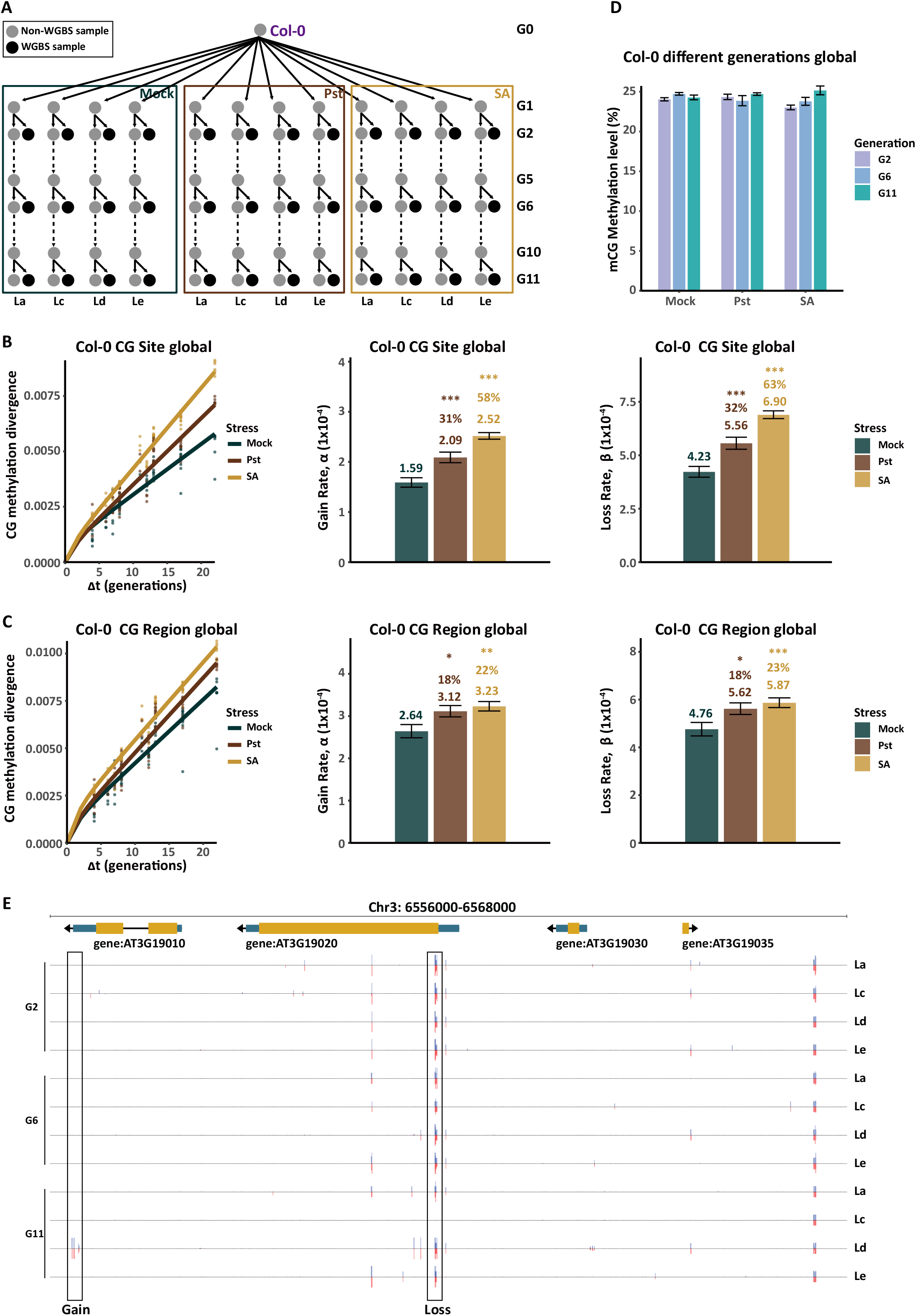
Multi-generational biotic stress increases the genome-wide epimutation rate. **(A)** Construction of Col-0 MA lines in the presence of two multi-generational biotic stressors: Pst and SA, and in the control condition (Mock). The MA lines in each treatment had 4 lineages(labeled as La, Lc, Ld, and Le) with WGBS samples collected in generation 2 (G2), 6 (G6), and 11 (G11). **(B)** Genome-wide (i.e. global) estimates of the per-CG epimutation rates (i.e. site-level) in the Mock, Pst, and SA conditions, separately. Left panel: CG methylation divergence plotted as a function of divergence time (Δt) for each treatment. Middle panel: estimated methylation gain rate (α) and the percent increase comparing the stress group and the control group. Right panel: estimated methylation loss rate (β). A two-sided t-test was used to test for rate differences between treatment groups. The asterisks indicate significant differences from Mock (* significant (0.01 ≤ P < 0.05); ** very significant (0.001 ≤ P < 0.01); *** highly significant (P < 0.001)). **(C)** Genome-wide (i.e. global) estimates of the per-CG region epimutation rate estimates. The notation is as in (B). **(D)** Steady-state genome-wide CG methylation levels are relatively constant across generations and are not significantly affected by the stress treatments.**(E)** Examples of spontaneous epimutations in the MA-lines. Gain: stochastic methylation gain; Loss: stochastic methylation loss.

Since CG methylation changes that occur in cytosine clusters are potentially more relevant functionally (23, 24), we also followed the methylation status of 100 bp regions through generations (region-level analysis). In this case, we defined an epimutation as a region-level methylation status change (14) (**Fig. 1E**). Region-level estimates revealed a similar pattern as seen in the site-level analysis, with multi-generational exposure to Pst and SA leading to an ∼18% and 23% epimutation rate increase, respectively (**Fig. 1C, Table S3**). As expected from theoretical models (12, 22, 25), the nearly proportional increase in both the gain and loss rates is consistent with CG methylation levels remaining largely unaltered across treatments and generations (**Fig. 1D**). This type of robustness in genome-wide steady-state methylation is a common observation in diverse stress studies in *A. thaliana* (6, 21, 26–35).

To compare our results with previous multi-generational abiotic stress experiments, we used the same analysis pipeline to reanalyze DNA methylation data of two MA lines grown under high salinity and drought conditions (6, 27). The salinity MA line spanned 13 generations with ∼5 WGBS per treatment, while the drought MA line covered 7 generations with ∼9 WGBS per treatment (**Fig. S1A-B**). We found that multi-generational exposure to salinity resulted in a 24% increase in the gain and a 27% increase in the CG methylation loss rates, respectively (**Fig. S2A, Table S5**)(6). A similar, albeit less pronounced, trend was detected in the drought treatment, although this increase was not statistically significant (**Table S5**)(27). This may be due to the drought treatment having been relatively mild compared to other stressors. Notably, global CG steady-state methylation levels remained relatively stable across different generations and treatments, regardless of whether the stress was biotic or abiotic (**Fig. 1D, Fig. S1C-E, Table S1, Table S3, Table S5**). Further analysis of the CG epimutation accumulation dynamics revealed no statistical support for selection at the genome-wide scale in any of the datasets, at least over the experimental timeframe investigated here (**Fig. 1, Fig. S2, Table S3, Table S5-S6**). Hence, stochastic gains and losses of DNA methylation appear to be effectively neutral under multi-generational stress, corroborating previous findings (12, 18, 22).

Taken together, our results demonstrate that long-term exposure to both biotic and abiotic stress compromises DNA methylation maintenance across generations, and leads to an accelerated accumulation of genome-wide spontaneous CG epimutations.

### Stress-induced epimutation rate changes are biased toward lowly methylated, transcriptionally non-responsive genes

In order to assess if stress-responsive epimutations accumulate more rapidly in specific genomic contexts, we estimated gain and loss rates separately in genes, promoters and transposable elements (TEs) (**Fig. 2A**) as well as in 36 annotated chromatin states (CS) (36, 37) (**Fig. S3**). While significant rate increases were detected in all genomic annotations, the largest increases occurred in genes (**Fig. 2A**) and their corresponding CS (**Fig. S4**). Consistent with the genome-wide analysis, these rate changes did not result in altered steady-state methylation levels in any of these annotations (**Fig. 2A**). Further partitioning of genes into those that are classified as body methylated (gbM) (N=5314, average CG steady-state methylation level = 31%), lowly methylated (LM) (N=12684, average CG steady-state methylation level = 2%) and all other genes (N=9452, average CG steady-state methylation level = 14%) revealed that the largest rate changes were concentrated in LM genes, while gbM genes were less affected (**Fig. 1E, Fig. 2B, Fig. S5, Table S3, Methods**). Indeed, the combined gain and loss rates in LM genes increased by an average of 26% under Pst and SA conditions relative to controls. This compares with an average change of 18.5% in gbM genes **(Table S3)**. These effects were not specific to biotic stress, but could be seen also under abiotic stress (**Fig. S2B-C, Table S5**), suggesting that they are a general stress signature.

**Fig. 2.**
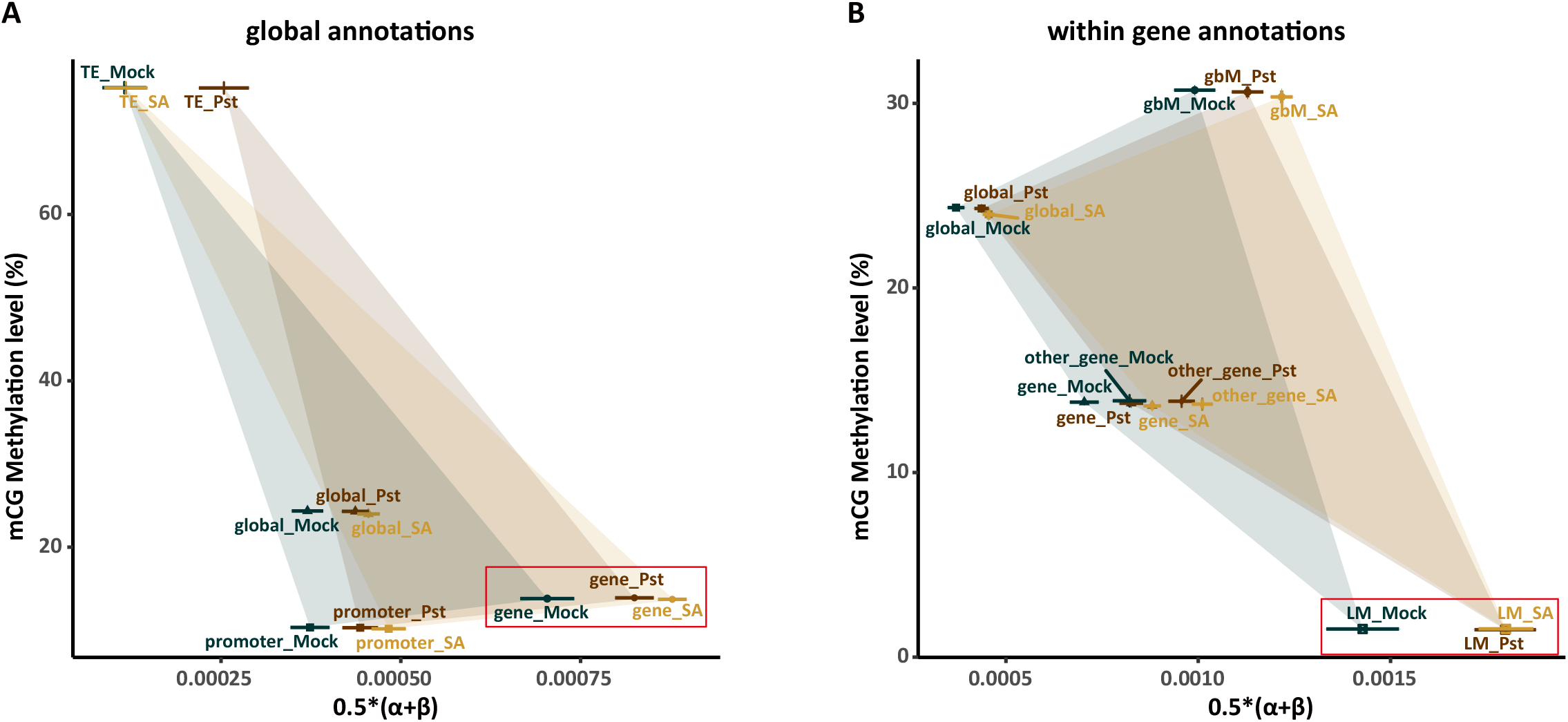
Hyper-accumulation of stress-responsive CG epimutations concentrates in LM genes. **(A)** Annotation-specific CG epimutation rate analysis in Col-0. The highest CG epimutation rate increase under Pst and SA stress were found in genes (x-axis). These rate increases occurred despite steady-state CG methylation levels remaining relatively constant within each of the annotations across treatments (y-axis). **(B)** Gene-specific CG epimutation rate analysis in Col-0. Genes were partitioned into gbM genes (N=5314), LM genes (N=12684), and other genes (N=9452) (see Methods). LM genes displayed the highest epimutation rate increases in Pst and SA relative to controls (x-axis). These rate increases occurred despite steady-state CG methylation levels remaining relatively constant within each of the gene annotations across treatments (y-axis).

Unlike gbM genes, which are conserved housekeeping genes (38) with low cell-to-cell or tissue-to-tissue transcriptional variation (39–41), LM genes are enriched for developmentally regulated and stress-inducible genes (**Table S7**). Gene Ontology (GO) enrichment analysis revealed that gbM genes are primarily associated with core biological processes such as DNA repair, while LM genes are mainly enriched in functions related to environmental stress responses, including hypoxia response (**Fig. S6A**). Further analysis of epigenetic marks showed significant differences in enrichment between LM and gbM genes. As expected, relative to gbM genes, LM genes were abundant in H3K27me3 (294%) and H2A.Z (123%) and depleted in H3K4me1 (47%) and H3K36me3 (44%)(**Fig. S6B**). Previous studies have shown that H3K27me3 levels are linked to plant stress responses (42–46), and that H2A.Z has a role in environmental sensing (46–48).

We therefore reasoned that the epimutation rate increases observed in LM genes are simply a byproduct of transcriptional responses to biotic stress. Active transcription might interfere with proper DNA methylation maintenance by both methylases and demethylases that can co-target these types of genes. To test this hypothesis, we re-analyzed transcriptome data from Halter et al. (49) (**Methods**). The authors studied how active demethylases regulate transcriptional immune reprogramming to enhance resistance to Pst. In line with our hypothesis, LM genes displayed the largest transcriptional changes in response to Pst, on average, and gbM genes the least (**Fig. 3A**). To tease apart these transcriptional changes in more detail, we partitioned the Col-0 transcription-response distribution into deciles, ranging from genes that are strongly down-regulated in Pst (decile 1) to genes that are strongly up-regulated (decile 10) (**Fig. 3B**).

**Fig. 3.**
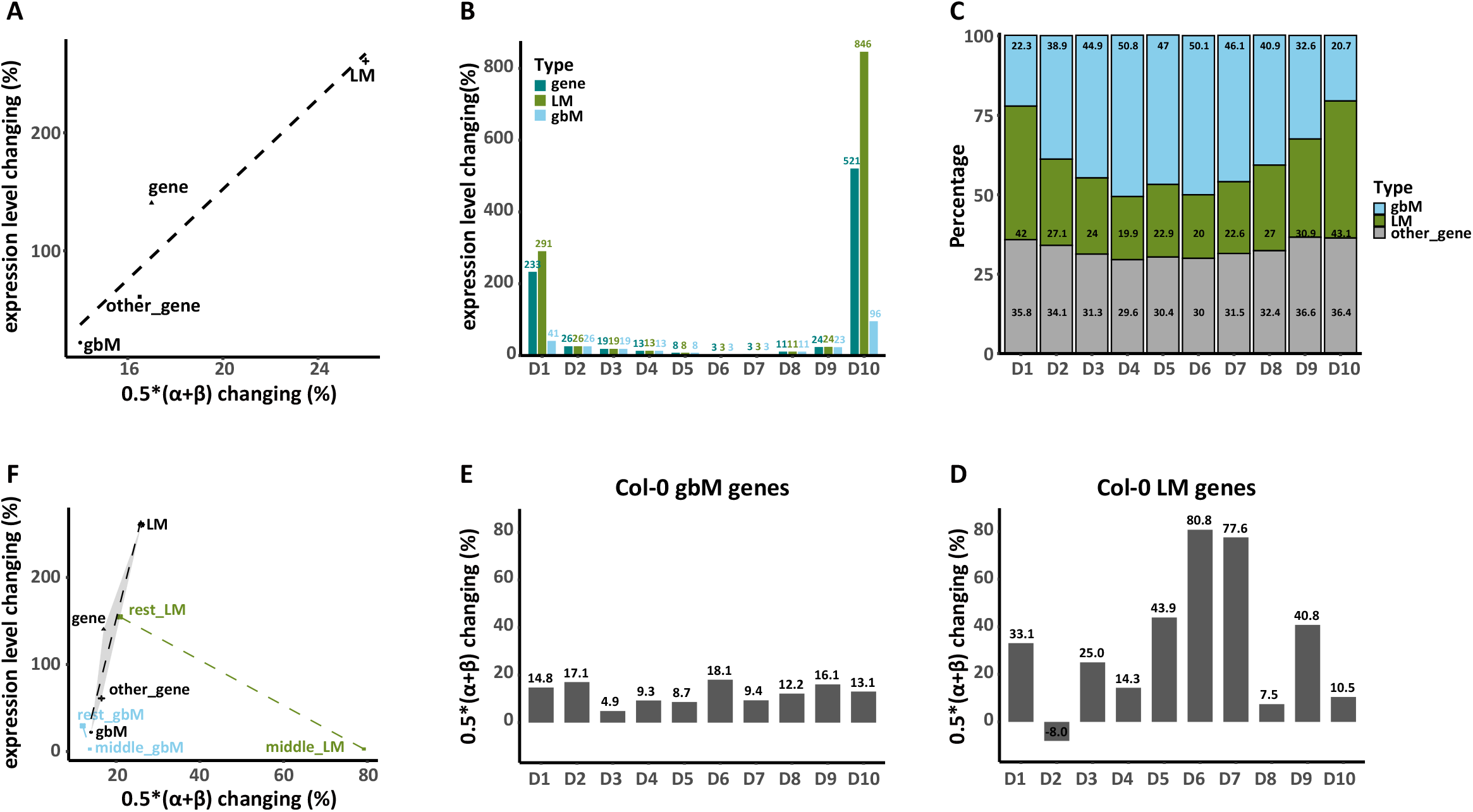
CG epimutation rate increases are most pronounced in LM genes that are transcriptionally non-responsive to stress. **(A)** Changes in expression levels (%) and CG epimutation rates (%) comparing Pst with Mock in Col-0. Genes were categorized gbM, LM, and other genes. The dashed line represents the linear fit, highlighting that LM genes show the most pronounced expression and epimutation rate changes. **(B)** Col-0 transcriptional response distribution to Pst: distribution of expression level changes (y-axis) of Pst relative to Mock, partitioned in 10 deciles (x-axis). Decile 1 represents genes that are strongly down-regulated under Pst and decile 10 represents genes that are strongly up-regulated. LM genes show the highest changes. **(C)** Stacked bar chart illustrating the percentage distribution of gene types (gbM, LM, other genes) across the 10 deciles of (B). LM genes are more prevalent in the deciles with high transcriptional responsiveness, indicating greater transcriptional changes, whereas gbM genes are concentrated in the deciles with low transcriptional responsiveness. **(D)** CG epimutation rate changes (in %) within Col-0 LM genes across the 10 deciles from (B). Deciles 6 and 7 exhibit the most significant changes, indicating that the largest rate increases occurred in a subclass of LM genes that were transcriptionally least affected by Pst exposure. **(E)** CG epimutation rate changes (in %) within Col-0 gbM genes across the 10 deciles from (B). Changes are more evenly distributed across all deciles. **(F)** Similar to (A), but including additional categories: “middle LM”, “middle gbM”, “rest LM”, and “rest gbM”. The notation “middle LM” and “middle gbM” represent the combination of deciles 6 and 7, while “rest LM” and “rest gbM” represent the combination of all other other deciles (i.e. excluding deciles 6 and 7). A subset of LM genes belonging to deciles 6 and 7 exhibit different patterns than the other deciles: they show the largest CG epimutation rate changes and the lowest expression level changes in response to Pst. By contrast, the pattern for gbM genes remains similar to (A), regardless of the deciles considered.

In line with our expectation, LM genes were most highly enriched in the extreme deciles, and gbM genes concentrated in the transcriptionally least responsive deciles (**Fig. 3C**). However, when we examined Pst-induced CG epimutation rate changes for each of the deciles separately, we found that the largest rate increases actually occurred in a subclass of LM genes that were transcriptionally least affected by Pst exposure (deciles 6 and 7, collectively referred to as “middle_LM”), while rate increases in gbM genes were approximately equally distributed among the ten deciles (**Fig. 3D-F, Fig. S7**). GO enrichment analysis revealed that the middle_LM genes are primarily enriched in metabolic processes, while rest_LM (representing all other deciles except 6 and 7) genes are more associated with salicylic acid and oxygen response pathways (**Fig. S8A**). A distinction between these two gene sets were also visible at the level of histone modification profiles, with rest_LM showing higher H3K27me3 (19%) and H2A.Z (10%) and lower H3K4me3(18%), H3K9ac(16%) and H3K36me3 (12%)(**Fig. S8B**). Hence, the functional differences between middle_LM and rest_LM is strongly associated with their differential epimutation rate response to stress.

### ROS1 mediates epimutation rate changes under multi-generational biotic stress

We sought to explore the connection between stress-induced transcriptional responses and epimutation accumulation in more detail. The DNA demethylase mutant *ros1* is a potentially informative experimental system for exploring this link. Recent studies demonstrated that ROS1 is crucial for the proper expression of genes during drought stress responses (50) and required for basal resistance to Pst (49). Re-analysis of transcriptome data from Halter et al. (49) revealed that *ros1* showed substantially altered expression responses to Pst, particularly in LM genes (**Fig. 4A**). LM genes that were most differentially expressed in Col-0 (deciles 1 and 10) were 3.24-fold less responsive to Pst in the *ros1* background, while genes that were not differentially expressed in Col-0 (deciles 6 and 7) showed a 4.41-fold induction (**Fig. 4B**).

**Fig. 4.**
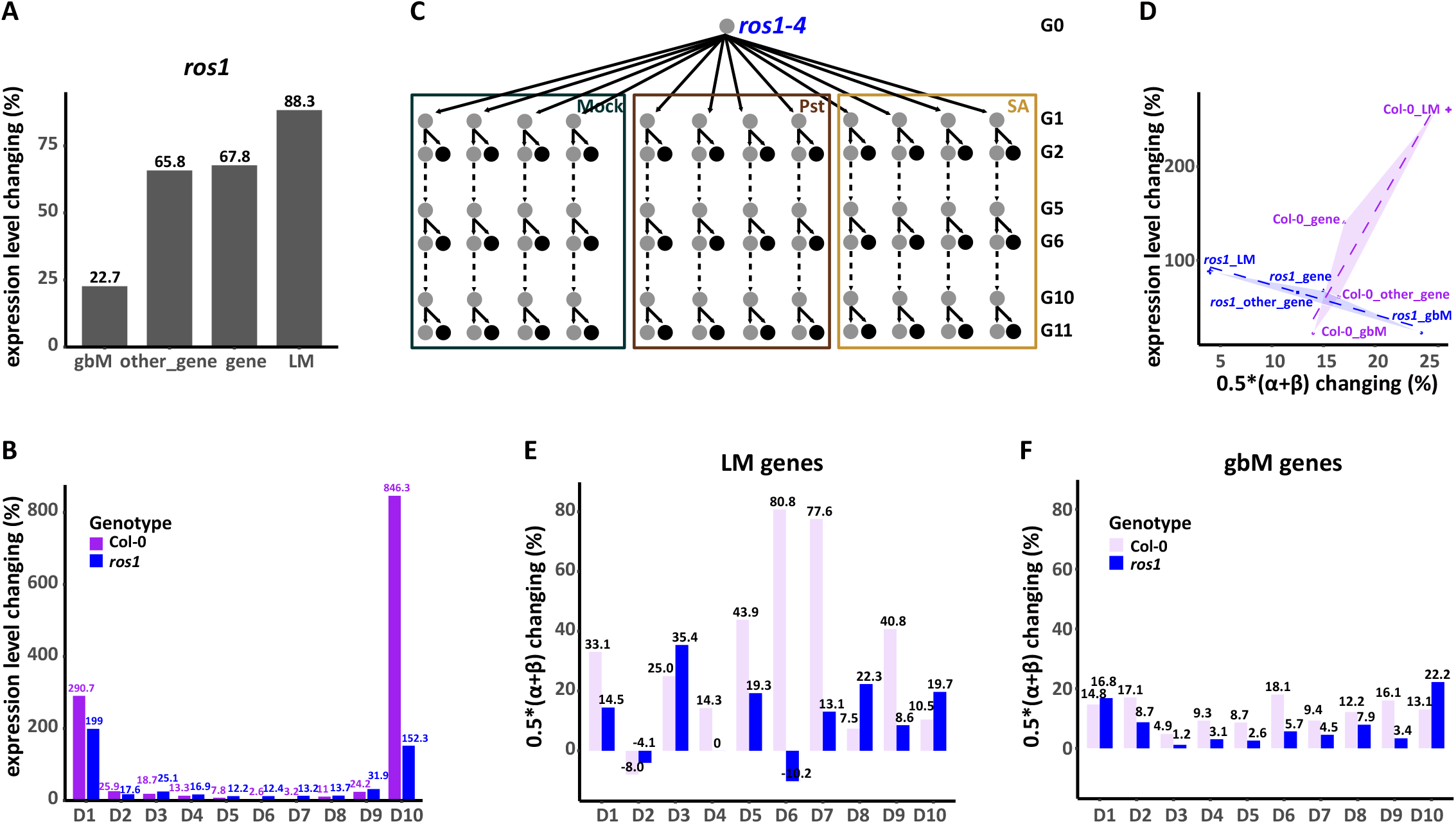
ROS1 mediates epimutation rate changes under multi-generational biotic stress. **(A)** Expression level changes between Pst and Mock in *ros1*, with genes categorized as gbM, LM, and other genes. **(B)** Distribution of expression level changes between Mock and Pst conditions for the 10 deciles for the Col-0 and *ros1* genotypes. **(C)** Similar to Col-0 in Fig. 1A, MA lines were constructed in *ros1-4* background under Mock, Pst, and SA conditions. **(D)** Similar to Fig. 3A, expression level changes and epimutation rate changes between Mock and Pst conditions for Col-0 and *ros1* genotypes. The dashed line represents the linear fit, highlighting the differences between the two genotypes. **(E)** The epimutation rate changes within LM genes across deciles for both Col-0 and *ros1* genotypes. Deciles 6 and 7 exhibit the most significant changes in Col-0 but are absent in *ros1*. **(F)** The epimutation rate changes within gbM genes across deciles for both Col-0 and *ros1* genotypes. Changes are evenly distributed across all deciles for both genotypes.

To assess the impact of these transcriptional changes on the epimutation landscape of *ros1* plants, we generated advanced-generation *ros1* MA lines under both control, SA, and Pst conditions in parallel, using the same experimental design as for Col-0 (**Fig. 4C, Table S8**). Consistent with two recent studies that analyzed *ros1* MA lines in ambient environments (13, 51), we found a 1.96-fold and 1.82-fold increase in the gain and loss rates at the genome-wide scale in our control condition compared to Col-0 (Gain rate: Col-0: 2.64×10^−4^ vs *ros1*: 5.18×10^−4^; Loss rate: Col-0: 4.76×10^−4^ vs *ros1*: 8.64×10^−4^)(**Fig. S9, Table S3, Table S9**), indicating that DNA methylation maintenance is generally compromised in this mutant. Specifically, under control conditions, LM genes were most affected by *ros1* in terms of epimutation rates and expression levels, while gbM genes were the least affected (**Fig. S9-S10, Table S3, Table S9**).

However, more striking was the fundamentally altered sensitivity of epimutation rates to *Pst* in this mutant background. Indeed, the stress-induced rate increases seen in Col-0, particularly in LM genes, were essentially absent in *ros1* (**Fig. 4D-F, Fig. S11**). This “buffering effect” was most pronounced for deciles 6 and 7 of the Col-0 transcription response distribution (i.e. middle_LM genes) (**Fig. 4E**). Epimutation rates in these deciles were essentially unchanged across treatments in *ros1* and not distinguishable from the other deciles, which contrasts the nearly 2-fold increase seen in Col-0 in these deciles. It is possible that the slight transcriptional induction of genes in deciles 6 and 7 in the *ros1* background was sufficient to stabilize CG methylation maintenance across environmental conditions by some unknown transcription-dependent mechanism. Another possibility is that genes in these deciles become preferentially hypermethylated in *ros1*, thus shifting the methylation steady-state in these genes. Recent work with DNA methylation mutants *cmt3* and *suv456* demonstrated that even slight shifts in steady-state methylation can result in increased epimutation gain and loss rates (36). However, patterns of hypermethylation in *ros1* were relatively uniform across annotation categories and deciles (**Fig. S9D, Fig. S12-S13**), and thus unlikely to mediate the differential patterns of epimutation rate changes seen in this mutant.

Despite remaining gaps in our mechanistic understanding, our work clearly highlights an important role of ROS1 in integrating environmental signals, not just at the transcriptional level, but also in terms of how CG methylation is maintained under multi-generational biotic stress. Whether these transcriptional changes drive altered epimutational processes or the other way around remains to be determined.

## Discussion

Our study demonstrates that multi-generational biotic stress increases the CG epimutation rate in *A. thaliana* (**Fig. 1B-C, Table S2-S3**). Contrary to the notion that these rate increases are biased toward genes that are transcriptionally responsive to stress, we found that they are actually concentrated in non-responsive genes. These genes are characterized by lower steady-state methylation levels (i.e. LM genes) and sparser CG density compared with body methylated (gbM) genes (**Fig. 3, Fig. S5**), whose epimutation rates remain most stable upon stress exposure (**Fig. 2-3**). One could hypothesize that it is these characteristics of LM genes that create particular problems for CG methylation maintenance upon environmental perturbations, as pre-existing methylation coupled with high CG density tends to ‘stabilize’ gains and loss dynamics (13, 52). However, this hypothesis also predicts that LM genes should appear as ‘epimutation hotspots’, even under control conditions; that is not the case (36).

Nonetheless, LM gene-biased epimutation rate changes under biotic stress do seem to have a direct (or indirect) link to the DNA methylation maintenance machinery. In *ros1* MA lines, where ROS1, a DNA demethylase, is lost, this effect is essentially abolished (**Fig. 4D-E**). How the loss of ROS1 accomplishes this is unclear (**Fig. 4**). We speculate that it is an indirect effect acting via *ros1*-induced transcriptional changes in response to stress. Indeed, loss of ROS1 leads to a subtle transcriptional induction of subsets of LM genes that were previously unresponsive in Col-0 (**Fig. 4B**), and - at the same time - buffers the transcriptional changes of subsets of genes that were previously stress-responsive in Col-0 (**Fig. 4D**). The net result is an overall contraction of the dynamic range of gene expression, making LM genes “look” more like gbM genes at the transcriptional level. It is therefore possible that constitutive, low/intermediate gene expression, which is characteristic of gbM genes (53), is a way to stabilize stochastic methylation gain and loss dynamics across environmental conditions.

Interestingly, recent work shows that gbM genes exhibit reduced transcriptional noise compared to non-gbM genes (39–41). This has led to the suggestion that the presence of CG methylation provides a way to stabilize gene expression states (39–41). This certainly makes sense, given that gbM genes have mainly housekeeping functions (38) and need to be tightly regulated across cells, tissues, and environments. This same reasoning predicts that LM genes - which are environmentally responsive (on average) - should exhibit high transcriptional noise, and that changes in their expression variability are somehow linked to DNA methylation maintenance fidelity under stress.

To explore this point, we generated RNA-seq data for 16 Col-0 plants in the Mock and SA treatments (**Fig. 5A, Fig. S14**), and quantified gene expression variability across plants within each of the treatments (**Methods**). We then partitioned the “noise” (i.e. variability) distribution into deciles, ranging from the least noisy (decile 2) to the noisiest (decile 10) genes (**Fig. 5B**). We found that gbM genes were enriched in lowest (decile 2) while LM genes were most prevalent in highest (decile 10) deciles of the noise distribution (**Fig. 5C-D**). This observation is consistent with our expectation as well as with recent findings by Zastąpiło et al. (40), who reported that genes with high variability in expression are enriched in stress response genes (i.e. our LM genes) based on GO analysis.

**Fig. 5.**
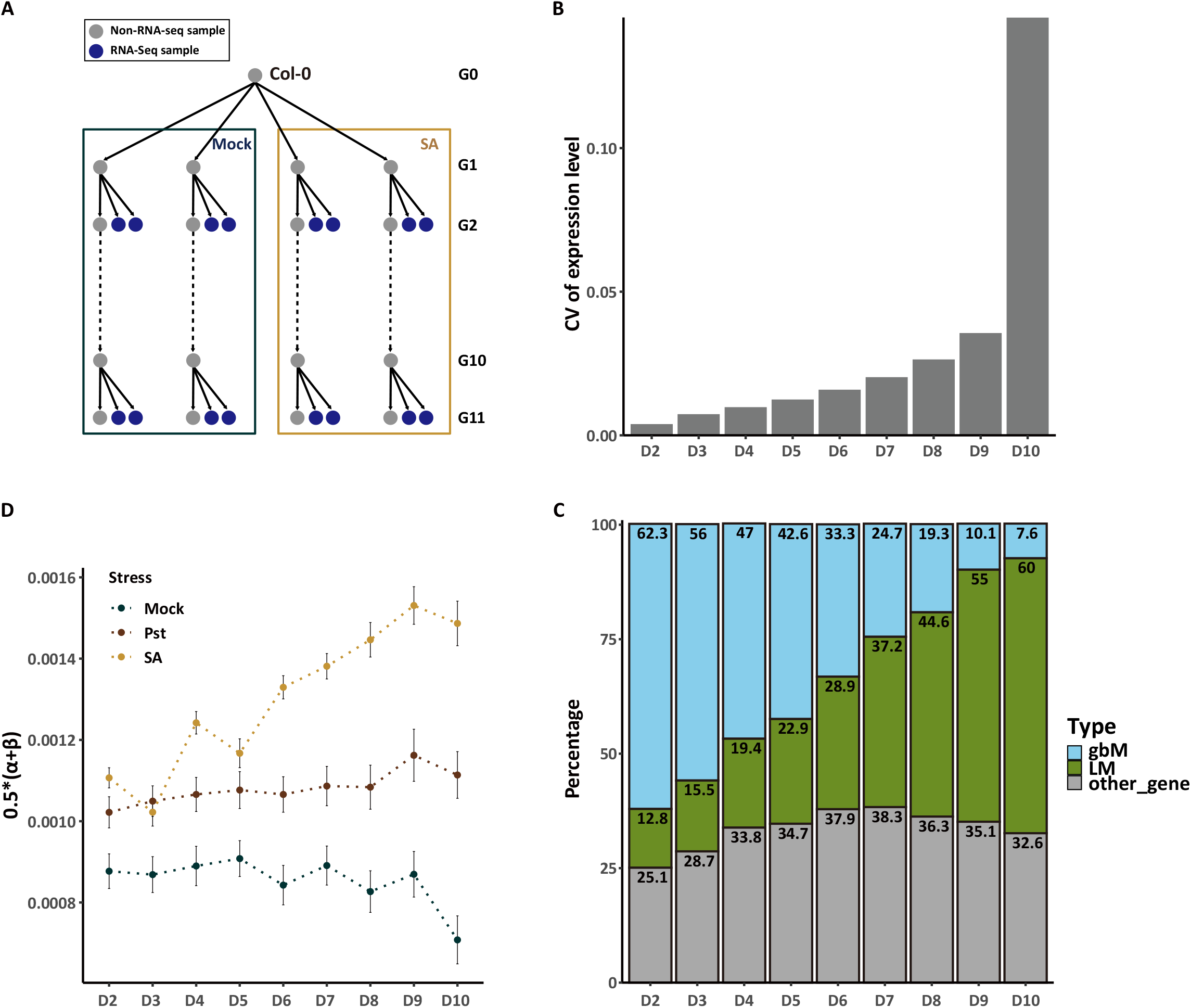
The rate increases are particularly pronounced in subsets of genes with high transcriptional noise. **(A)** Overview of RNA-seq sampling strategy under multi-generational biotic stress with SA and a control condition (Mock). RNA-seq samples were collected from two lineages in both Mock and SA treatments at generation 2 (G2) and generation 11 (G11). **(B)** According to the Coeffcient of Variation (CV) of expression level in the same gene among different samples in Mock, we discriminate 10 gene subsets according to their quantiles. The CV of expression level is increased from the D2 gene set to the D10 gene set gradually. D1, representing constitutively unexpressed genes, was removed to focus on expressed genes. **(C)** From the D2 gene set to the D10 gene set, the percentages of LM genes increase and the percentages of gbM genes decrease gradually. **(D)** From the D2 gene set to the D10 gene set, the epimutation rate difference between the control group (Mock) and the stress group (Pst and SA) increases gradually. Combining with B and C, they show that stress-responsive rate increases are particularly pronounced in subsets of genes with high transcription noise, which are mostly LM genes.

We suspect that the transcription-noise profile in a *ros1* background is substantially altered, indicative of a dysregulated transcriptome. In fact, studies have shown that *ros1* mutants display reduced resistance to pathogens, which may reflect a decreased ability of their transcriptome to cope with stress (49, 54, 55). ROS1 target sites are associated with changes in the levels of histone modifications, including H3K27me3, H3K4me3, H3K36me3, and H3K9ac (56). Furthermore, H2A.Z physically interacts with ROS1 to regulate DNA demethylation and anti-silencing (57). Mutations in ROS1 disrupt the H2A.Z-dependent DNA demethylation process and may be sufficient to facilitate the exclusion of histone variant H2A.Z, which has a role in environmental-sensing (47, 48) and acts antagonistically with DNA methylation (58). Hence, changes in the chromatin-environment linked to H2A.Z density within gene bodies may mediate both the transcriptional as well as the epimutation rate changes observed in our study. A detailed mechanistic insight into this process warrants further investigation.

Our work has highlighted a subset of LM genes as major sites of epimutation rate changes under stress. Since we could show that these genes are transcriptionally non-responsive to biotic stress in Col-0, we suspect that CG epimutations that accumulate in them are effectively neutral. By extension, one could conclude that CG epimutational processes under stress have no bearing on long-term adaptive processes in plants. However, we note that this conclusion is based on averages over many genes. We therefore cannot rule out that CG epimutations occurring in other subsets of LM genes that are stress-responsive could be functionally important, even if these events occur at very low frequencies. In fact, we identified some examples of LM genes that contained > 30 mCG (**Fig. S15A**), even though they were previously identified as unmethylated in other datasets (62). This number of mCG is similar to what is typically found in gbM genes, on average (**Fig. S15B**). It indicates that LM genes have the capacity to be stochastically targeted for larger methylation changes, which may, or may not, be functionally relevant. The acquisition of these latter methylation changes could lead to segregating epialleles in natural populations with potentially long-term evolutionary consequences. The stability and fitness effects of such putative epialleles would need to be assessed on a case-by-case basis in future studies.

## Materials and Methods

### MA line construction and plant material

MA lines of both Col-0 and *ros1-4* genotypes were propagated with four lineages per treatment from the same ancestor by single seed descent for 12 generations. Plants from G1 to G11 were recurrently exposed to *Pseudomonas syringae* or salicylic acid, which was applied to leaves at 3, 4, and 5 weeks of age, except for the sequenced samples. Samples were sequenced at generation 2 (G2), generation 6 (G6), and generation 11 (G11) using a sibling design (22) (**Fig. 1A, Fig. 4C**). In details, Arabidopsis thaliana seeds were suspended in 0.1% agarose (0.1% w/v agarose in deionised H2O (dH2O)) and stored in the dark for 2 days at 4 °C to break dormancy, after which they were sown onto soil consisting of Levington Advance Pot & Bedding M3 compost (ICL) and sand in a 4:1 ratio and cultivated under the following conditions: 16:8 hr day: night, 22 °C, 60% relative humidity (RH) and 150 μmol m−2 s−1. At 10-14 days, seedlings were thinned to one per pot. Lines were propagated through collection of seed from single plants. At 21, 28, and 35 days post-sowing, plants were treated with a mock treatment, salicylic acid (SA), or a pathogen. 10 mM MgSO4 was used as the mock treatment, while salicylic acid was delivered as 5 mM sodium salicylate in 10 mM MgSO4. For the pathogen treatment, 100 mL LB culture of *Pseudomonas syringae* pv. *tomato* DC3000 (Pst) (59) was grown at 25-28ºC with shaking at 200 rpm. The Pst was resuspended in 10 mM MgSO4 at OD 0.1 (595nm). 0.01% Silwet L-77 was added to each solution immediately before plant spraying. Plants were covered for two days after each treatment. Seedlings were cultivated under the same growth conditions. After 3 weeks of growth, 28-35 seedlings per replicate were extracted, pooled, flash frozen and ground to a powder. Genomic DNA was extracted by Leonardo Furci using the GenElute Plant Genomic DNA Miniprep Kit (Sigma-Aldrich).

The experimental design for the two published MA lines that were grown under multi-generational abiotic stress (saline and drought stress) is summarized in **Fig. S1A-B**. Further details concerning the construction of these pedigrees and DNA extraction protocols can be found in their original publications (6, 27).

### WGBS and data processing

Sequenced using 150 bp paired-end reads on the Illumina platform. All WGBS data were processed using the MethylStar pipeline (22, 60) with TAIR10 (61) as the reference. Different files corresponding to the same sample were merged at the BAM file stage using “samtools merge -n” (Samtools version 1.11). For all samples, Methimpute (18, 60) was used to call three cytosine-level methylation states: unmethylated (“U”), intermediate methylated (“I”), and methylated (“M”). To ensure high-quality methylation state calls, cytosines with a maximum posterior probability larger than 0.99 were retained. The sequencing depth, mapping efficiency, coverage, bisulfite conversion rate, total reads number and unique reads number are detailed in **Table S1 and S8**.

### RNA-Seq and data processing

Leaf RNA-seq samples were acquired from the siblings of progenitors in two lineages under Mock and SA treatment in both G2 and G11 in the Col-0 genotype. Similar to the WGBS sampling design, the plants used for RNA-seq sequencing in the SA treatment were not exposed to biotic stress, although their parents were. The treatments were applied as foliar sprays on days 21, 28, and 35 after sowing (DAS). The mock treatment consisted of 10 mM MgSO4, while the SA treatment involved 5 mM sodium salicylate dissolved in 10 mM MgSO4. Both solutions were supplemented with 0.01% Silwet L-77 as a surfactant. After these treatments, the G1 plants were allowed to set seeds, producing the G2 generation. This treatment cycle was repeated for nine more generations, resulting in a G11 seed stock. 16 two-three week old seedlings (∼6-8 true leaves) per treatment were snap frozen in liquid nitrogen and total RNA was extracted using Trizol. Sequenced using 150 bp paired-end reads on the Illumina platform.

We also reanalyzed previously published RNA-seq data of Col-0 and *ros1* under Pst treatment (49). In Halter’s data (49), the so-called Pst treatment actually involved exposure to flg22, and the specific *ros1* mutant analyzed was *ros1-3*. This allowed us to study the direct effects of biotic stress on gene expression and to explore the relationship between expression levels and epimutation rates in conjunction with our WGBS data.

For our RNA-seq samples, reads were trimmed using Trimmomatic (version 0.38) and aligned with STAR (version 2.7.6a). Counts were gathered using HTSeq-count (version 0.13.5). The analysis was conducted using TAIR10 (61) as the reference genome along with the Araport11 annotation. Differential expression analysis was performed using DESeq2 (version 1.30.1), which included the removal of batch effects in our RNA-seq data. For both our RNA-seq samples and Halter’s published RNA-seq data, rlog normalization was applied separately to each dataset. After performing rlog normalization on Halter’s RNA-seq data for both Col-0 and ROS1 genotypes, genes with expression levels below 1.5 in both the control and treatment conditions were excluded from the analysis to focus on expressed genes. Subsequently, to analyze Halter’s RNA-seq data, we categorized the genes into ten deciles (named by D1 to D10) based on the ranked percentage changes in expression levels between control and treatment conditions in the Col-0 genotype.

For the definition of transcriptional noise, we calculated the Coefficient of Variation (CV) of expression levels for each gene across different samples under Mock conditions in our RNA-seq data. The CV for a given gene is defined as follows: CV = (standard deviation of samples) / (mean of samples). Based on CV ranking, we identified ten gene deciles named D1 to D10 for transcriptional noise analysis. D1, representing constitutively unexpressed genes, was removed to focus on expressed genes.

### Enrichment of annotations

Annotation files for genes and transposable elements (TEs) in GFF3 format were downloaded from Ensembl Plants (http://plants.ensembl.org/info/data/ftp). Promoters were defined as regions 1.5 kb upstream of the transcription start site (TSS). The lists of gbM and LM genes were obtained from previous work (62). Using a per-gen binomial approach, the authors classified gbM genes as those with significant CG methylation, requiring at least 20 CG sites with a q-value < 0.05 for mCG, and q-values > 0.05 for mCHG and mCHH. LM genes were defined by having read coverage of at least 20 CHH sites with q-values > 0.05 for all contexts (CG, CHG, CHH) and less than 2 symmetrical mCG sites. We note that previous work has referred to LM genes as UM genes (i.e. unmethylated genes) to draw a clear contrast with gbM genes (62). However, as can be seen in **Fig. 1E, Fig. S5, Fig. S15, Table S3, and Table S9**, these genes feature low steady-state CG methylation, on average, and the classification of specific genes containing only unmethylated CGs is relatively fluid across lineages and generations (**Fig. S16**). Finally, we defined “Other_gene” as the remaining set, after removing gbM and LM genes from the entire gene set. “Other_gene” includes teM genes, which are defined by significant CHG or CHH methylation, with at least 20 mapped sites and a q-value <0.05 for CHG or CHH methylation. The individual epigenetic marks and genome coordinates for the 36 CSs were obtained from the PCSD database (37). Following the method of Hazarika et al. (36), we clustered the 36 CSs into four broad groups, represented by green, red, blue, and purple states.

### Estimation of epimutation rates

Following Hazarika et al. (36), global and specific annotated estimates of the CG methylation gain rate (α) and the loss rate (β) were acquired using the R package AlphaBeta (version 1.10.0)(22). A neutral model (ABneutral) was fitted in all cases. Two selection models (ABselectMM and ABselectUU) were tested on global epimutation rates across all treatments, including biotic stress and abiotic stress.

### Calculation of average methylation levels

Average methylation levels were calculated with posteriorMax ≥ 0.99 for global and specific annotations.

